# GeneScribe - a tool for fact extraction and gene summarization using Large Language Models (LLMs)

**DOI:** 10.64898/2026.11.25.769256

**Authors:** Pengyuan Li, Ranjana Kishore, Valerio Arnaboldi, Paul Sternberg

## Abstract

The rapid growth of biological data presents major challenges for expert curation and integration of gene information into biological knowledgebases such as the Alliance of Genome Resources (Alliance), limiting data discovery, analysis and dissemination. The current automated approach to gene summary generation at the Alliance relies on curated data and a combination of rule-based and optimization algorithms. While effective, this method is limited by the timely availability of curated annotations, and prone to producing redundant or fragmented descriptions that may obscure the key biological roles of a gene. To address these limitations, we developed GeneScribe, a tool that leverages Large Language Models (LLMs) to automate and scale the generation of gene function summaries. GeneScribe employs a two-step workflow: (i) automated fact extraction from the scientific literature, producing gene-specific statements with related provenance without needing manual curation (ii) synthesis of concise, modular and rigorous summaries of gene function from the extracted facts. GeneScribe can provide automated updates to genes with existing summaries and generate timely new summaries directly from the literature for genes that lack one. This will reduce latency in including emerging findings while broadening genomic coverage of the automated gene summaries at the Alliance.

## Introduction

A textual gene summary provides a concise and easily accessible overview of key information about a gene, such as its molecular function/identity, participation in biological processes and relevance to disease, offering a single, unified view of critical gene-related information in one place. A gene summary is easy to read and does not require specialized knowledge of biological ontologies, controlled vocabularies, or familiarity with knowledgebases. However, generating and maintaining manually written gene summaries is labor- and resource-intensive (Kishore et, al. 2020). In particular, with the rapid accumulation of genetic and genomic data and limited curation resources, it is difficult to keep summaries current. Even though nowadays AI can generate gene summaries, using generic LLMs results in summaries that are not standardized and often not entirely based on available high quality curated data. Moreover, providing gene summaries in knowledgebases makes them reusable and more cost efficient.

In previous work at the Alliance of Genome Resources, a knowledgebase for the genetics and genomics of the major model organism species (Alliance; www.alliancegenome.org; Alliance of Genome Resources Consortium 2024), we implemented a heuristic-based optimization algorithm for automated gene summary generation. The algorithm combines rule-based methods and predefined sentence templates to generate summaries from structured, curated data (Kishore et, al. 2020). The optimization step of the algorithm balances information content with readability, which results in accurate summaries for curated genes. However, this approach is limited by the availability of curated data of new gene-related literature. For example, for *C. elegans*, while summaries are available for a substantial portion of protein-coding genes included in the Alliance, 25% of these genes (4,999 out of 19,983) lack any summary. For non–protein-coding genes—including non-coding RNAs and pseudogenes—the coverage is even more limited, with only 295 of 28,795 having any form of summary. This reveals a substantial gap in curated gene knowledge and highlights the limitations of approaches that are based on curated data only. LLMs offer a promising avenue to enhance gene summary coverage and comprehensiveness. In fact, a LLM-based approach can be used to automatically retrieve data from newly published literature, and generate new gene summaries as well as integrate new data into existing ones.

Here we present GeneScribe, a novel tool that uses LLMs to generate gene summaries at the Alliance with a two-stage workflow: (i) structured prompt-based fact extraction that converts unstructured text from publications into standardized gene-fact pairs and (ii) gene summary generation, where extracted facts are synthesized into concise, accurate, and stylistically consistent gene summaries. By identifying and extracting text from the paper as evidence for the fact extraction, GeneScribe improves transparency, reduces hallucination risk by using strictly defined prompts for both the fact extraction and generation phases, and enables independent validation of extracted statements prior to summary generation. Specifically, each extracted gene fact is linked to supporting text from the source publication, providing provenance. GeneScribe differentiates between two key gene groups: (i) genes that lack summaries at the Alliance and (ii) genes with existing summaries in the Alliance database that need updating. For the first group, GeneScribe generates gene summaries directly from the extracted facts. For the second group, GeneScribe uses a context-aware approach to retrieve the existing summary from the Alliance database and then updates it by incorporating new facts extracted from recent publications. This dual approach ensures that summaries use manually curated data when available, while including the latest available information from recent publications not yet curated. Importantly, this design enables summary generation for genes with limited or no curated annotations, thereby extending coverage beyond the constraints of the current gene summary generation system at the Alliance. The methodology described in this paper can also be easily integrated into the existing agentic AI solutions at the Alliance as its workflow is modular and already uses data sources and controls from the Alliance APIs and tools (Arnaboldi et al., 2026). A summary of the GeneScribe workflow is presented in Figure 1.

**Figure 1.**
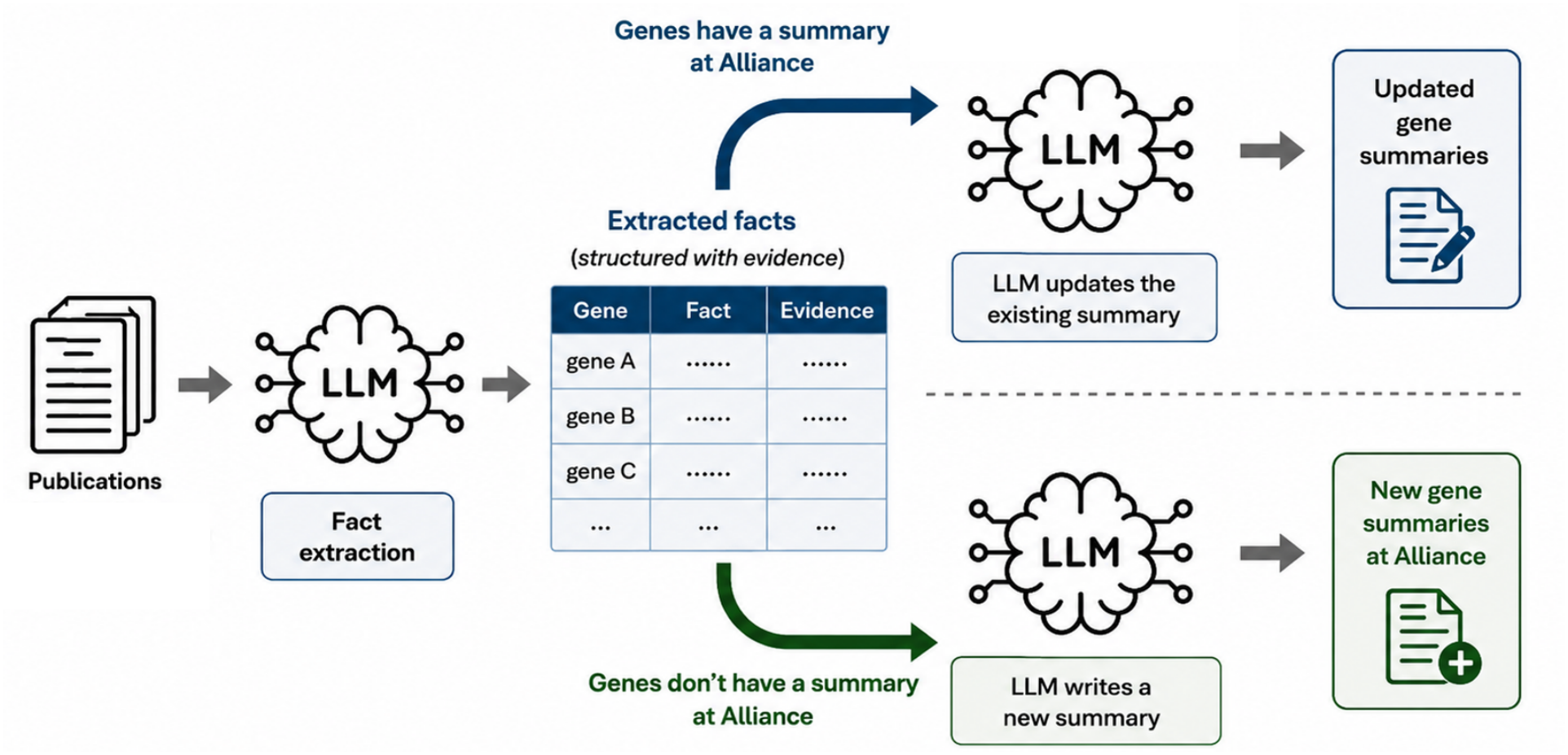
Gene summary generation workflow

Recent studies have increasingly explored the use of LLMs for gene-related knowledge question answering, functional interpretation, and biological reasoning. GeneGPT (Jin et al., 2024) augments LLMs with NCBI Web APIs to answer genomics questions involving gene nomenclature, genomic location, gene–disease associations, functional information, and multi-step reasoning. In contrast, GP-GPT (Lyu et al., 2025) develops genomics-specialized LLMs trained on structured and unstructured genomics and medical genetics data to support gene–phenotype knowledge representation, information retrieval, and relationship analysis. At the gene-set level, LLMs have also been applied to infer shared biological functions from groups of genes and generate supporting explanatory analyses, demonstrating their potential for functional interpretation of omics-derived gene sets (Hu et al., 2025). Together, these studies demonstrate the potential of LLMs to retrieve, reason over, and synthesize gene-related biological knowledge.

Several other methods for automated gene and biomedical summary generation have been proposed in the literature. For example, LitSumm (Green et al. 2025) applies LLMs to the summarization of scientific publications, with a focus on noncoding RNAs, demonstrating that domain-adapted models can effectively capture salient biological information from unstructured text. Similarly, Gene Set Summarization using LLMs (Joachimiak et al. 2024) has explored the use of generative models to produce concise descriptions for groups of genes, showcasing the potential of LLMs to synthesize heterogeneous information into coherent summaries. Most closely related to our work is GENESUM (Chen et al. 2024), an LLM-based framework for gene summary extraction. GENESUM employs a two-step workflow: (i) filtering sentences from the literature by keyword frequency and semantic similarity to ontology terms, and (ii) summarizing the filtered content using a fine-tuned LLM. While effective, this approach depends heavily on predefined filters and ontology-based similarity measures, which may bias the input set and exclude potentially relevant findings that fall outside these constraints.

Moreover, many LLM-based summarization approaches do not track individual statements back to specific evidence. Compared to existing solutions, GeneScribe preserves provenance for each extracted fact, allowing downstream applications to reuse them for sustainable integration and cost efficiency, facilitating manual validation and the retrieval of evidence data. Moreover, GeneScribe follows strict stylistic and scientific rules, making the generated summaries more harmonized and comparable between each other, which is particularly important for the Alliance.

## Materials and Methods

In order to test and validate GeneScribe, we collected a sample of species-specific scientific publications related to *C. elegans* from the Alliance literature store (Alliance Bibliography Central; ABC). GeneScribe used this sample to extract gene facts and then generate gene summaries for all the genes identified in the publications. Further, we performed an analysis on the performances of the extracted facts and generated gene summaries with the help of a professional biocurator.

### Source of Publications and Selection Criteria

For the analysis reported here, we chose to test GeneScribe on a set of *C. elegans* papers, but the same methodology can be extended to other organisms included in the Alliance. The ABC serves as the centralized platform for literature triaging and management of literature relevant to Alliance species, including the *C. elegans* literature. The ABC selects publications from several sources including PMC and PubMed, also ensuring species and biological relevance of these papers through manual oversight by professional biocurators.

We generated a list of *C. elegans* genes that lack an automated summary at the Alliance but may have publications that contain data that have not yet been curated. To identify this target list of publications, we queried the ABC to obtain papers that mention any gene in our list in the abstract and/or title –to try to exclude papers with tangential gene mentions. The resulting publications likely contain uncurated data related to genes that lack a summary at the Alliance, but they may also contain novel data for genes that already have a summary at the Alliance. Thus, the sample we collected can be used for testing how well GeneScribe generates both de novo summaries and updates for existing summaries. To focus on experimental studies, we further filtered the ABC search to include only references classified as ‘journal articles,’ excluding reviews, conference proceedings, book chapters, and other publication types. Full-text documents were obtained in PDF format from the ABC and parsed into text using GROBID (GROBID, 2008), a machine learning tool that converts unstructured text from scientific publications into structured information that can be easily read by LLMs or converted to plain text. A total number of 83 publications were collected from the ABC search. These papers contained mentions to 212 genes, 156 genes with an existing summary at the Alliance and 56 without one.

### Steps in Gene Summary Generation

GeneScribe generates gene summaries using a two stage workflow. In the first stage, the system automatically extracts gene-specific facts from the literature, constrained by clearly defined biological data types and detailed prompts. In the second stage, these structured facts are synthesized into concise and biologically meaningful gene summaries through prompt-guided text generation.

### Step 1: Fact extraction

#### Data types chosen for fact extraction

Gene summaries generated by GeneScribe include a set of selected and well-defined datatypes from standards developed by the biocuration community and projects such as the Gene Ontology (GO) (Ashburner et al. 2000;The Gene Ontology Consortium 2026). These include:

##### Biological Process

Cellular and organismal processes in which the gene is involved.

##### Molecular Function

Specific biochemical activities performed by the gene product.

##### Gene Interaction

Physical or genetic interactions with other genes.

##### Pathway

Biological pathways that the gene is part of.

##### Gene Expression

Tissue and sub-cellular localization of the gene.

##### Relevance to Disease

Associations with human diseases.

These data types ensure that gene summaries contain a well-rounded representation of the gene’s biological significance, integrating functional and disease-related aspects.

These data types were selected to align with existing Alliance gene summary data types while extending coverage to data types such as interactions and pathways that were not previously included in the Alliance automated summaries.

#### Prompt Design

To extract relevant biological facts from scientific publications, GeneScribe employs a structured prompting framework that guides a LLM to generate outputs in a predefined, machine-readable format. This design ensures that all extracted facts are consistent, interpretable, and compatible with biocuration workflows.

The prompt is instantiated for each publication using three inputs: (i) the full text of the paper, converted from PDF into plain text format, (ii) a list of target genes identified from the title and abstract of the paper and (iii) a predefined JSON schema describing the expected output structure (see below). Rather than asking the model to summarize the paper directly, the prompt frames the task as structured fact extraction, requiring the model to identify gene function related to the specified data types. By limiting extraction to a set of target genes and requiring structured outputs, the prompt reduces noise from incidental gene mentions and encourages consistent extraction across publications.

The JSON schema defines a structured output format in which each gene is associated with a set of data type-specific properties (see above for the list of data types). To make extracted facts human readable and easier to validate, the schema also contains standardized sentence templates for each data type. The schema also requires the inclusion of one or more verbatim text spans from the source publication supporting each extracted fact. We refer to these lines of text as “evidence sentences”. Each gene-fact pair is therefore stored together with its supporting evidence, establishing explicit provenance between the structured assertion and the source publication. This design enables curators to inspect the original textual evidence when validating an extracted fact and allows the structured facts to be reused in downstream curation workflows without losing their connection to the literature.

A summary of the data type-level prompting strategy is provided in Table 1, and the complete extraction prompt, including the full JSON schema specification, is available at our GitHub repository (See Data Availability section). As an example, for the Biological Process data type, the prompt instructs the model to distinguish between experimentally supported findings and predictions using different sentence templates. For instance, experimentally supported statements are represented using the templates:

**Table 1.**
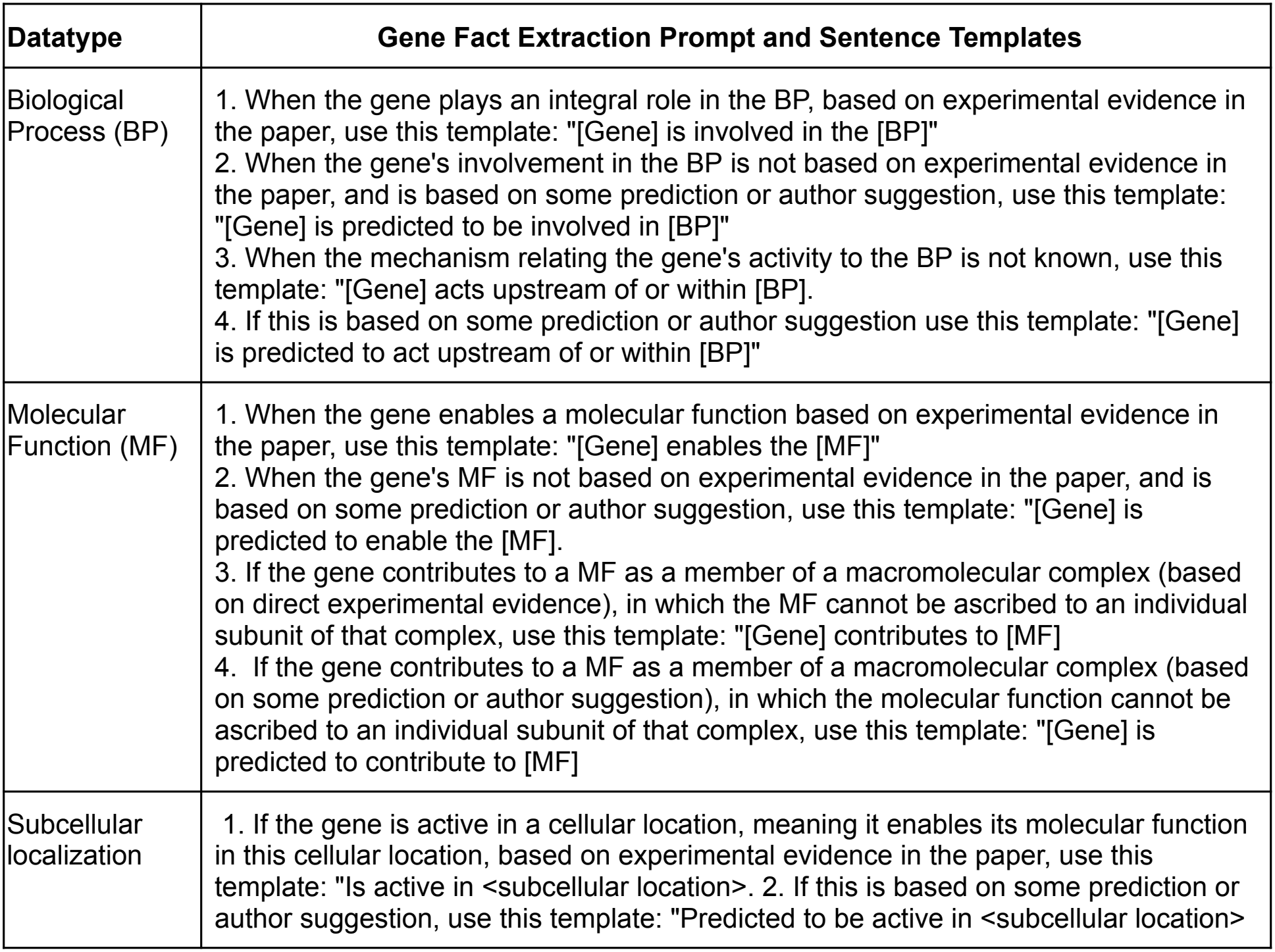
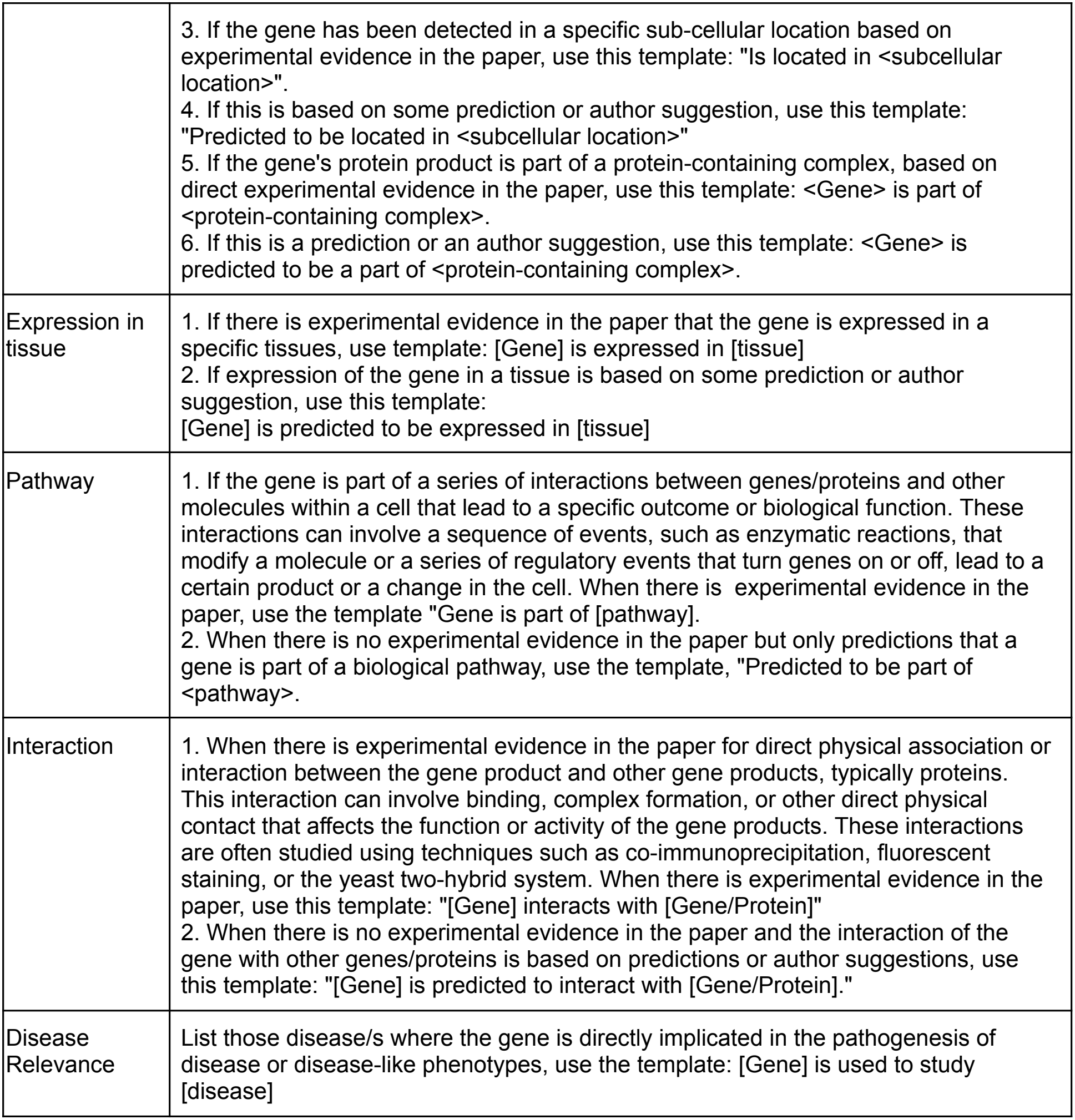
Instructions provided to the LLM with community standard definitions of the data types. Sentence templates for each data type are also provided.

*“[Gene] is involved in [Biological Process]”*

whereas predicted findings are represented *as:*

*“[Gene] is predicted to be involved in [Biological Process]*.*”*

Similar data type-specific templates are defined for molecular function, pathway, interaction, expression, localization, and disease relevance.

#### Models

To identify the most suitable model for the fact-extraction task, we evaluated several LLMs with comparable prompting configurations, including GPT-4-turbo, GPT-4o, GPT-5, and GPT-o1 (Achiam et al. 2023). Each model was assessed for its ability to generate structured, schema-compliant outputs that captured relevant biological information from the literature. Although all models demonstrated basic capability in extracting gene-level facts, their performance varied in terms of completeness, adherence to the JSON schema, and consistency in providing supporting evidence sentences. Among these, the GPT-o1 model produced the most comprehensive and well-structured results, consistently populating all relevant data fields and maintaining the required output format. Consequently, GPT-o1 was selected as the primary model for fact extraction and gene summary generation in GeneScribe. As this was a pilot study, we prioritized extraction performance over inference cost. At the time of the study, GPT-o1 had a higher cost per token compared to GPT-4o. Moreover, GPT-o1 is a reasoning model that uses additional reasoning tokens during inference, further increasing the overall inference cost. In future work, we will consider lower-cost and open-source models that will be more cost-effective.

### Step 2: Gene summary generation

Genes in our publication sample obtained from the Alliance were categorized into two groups (i) Genes that lack a summary (ii) Genes that have a summary at the Alliance that require updating based on new publications.

### Genes that lack a gene summary

For genes that lack a gene summary at the Alliance, GeneScribe generates a de novo gene summary directly from the structured facts extracted in Step 1. The prompt receives the extracted gene facts, organized by biological data type (e.g., molecular function, biological process, pathways, interactions, expression, and disease relevance), and instructs the LLM to synthesize them into a concise, coherent, and accurate description.

#### Prompt

The prompt for generating a gene summary for genes that lack one consists of four components: (i) **a task description** defining the objective of generating an Alliance-style gene summary from structured facts; (ii) **an example gene summary** that serves as an in-context demonstration of the desired writing style and organization; (iii) **data type-specific sentence templates** that were used in the fact extraction step to specify how different biological data types should be expressed and (iv) **summarization guidelines** that instruct the model to organize related biological facts into higher-level biological concepts, preserve the distinction between experimentally supported findings and predictions, omit data types with no available information, avoid redundant statements, and maintain a concise scientific writing style. Together, these prompt components encourage the model to generate standardized summaries that are stylistically consistent with existing Alliance gene summaries while accurately reflecting the structured information extracted from the literature.

### Genes with a gene summary

For genes with an existing Alliance summary, GeneScribe uses a context-aware updating strategy. This process consists of two steps: (i) Retrieval of existing summaries: the current Alliance gene summary is retrieved and (ii) Generation of an updated summary: the LLM integrates newly extracted facts into the existing summary.

#### Prompt

The prompt consists of four components: (i) **a task description** defining the objective of updating the existing summary (ii) **the current Alliance gene summary**, which serves as the primary context for revision (iii) the newly extracted structured facts organized by biological data type and (iv) **data type-specific sentence templates** and updating guidelines. The guidelines instruct the model to preserve existing information, incorporate newly extracted facts into the appropriate data types, summarize related ontology terms into higher-level biological concepts, remove redundant or repetitive information, and maintain the writing style and sentence structure used by existing Alliance summaries. Similar to the de novo summary generation prompt, datatype-specific templates (e.g., *“Enables*…*”, “Involved in*…*”*, and *“Expressed in*…*”*) are provided to ensure stylistic consistency across updated summaries. This strategy allows summaries to evolve systematically as new data becomes available while preserving the integrity of curated knowledge. In this way, GeneScribe supports scalable, accurate, and traceable updates to gene summaries.

### Evaluation

To evaluate the performance of GeneScribe, we checked both the fact extraction and the gene summaries using expert curator assessment. For the gene fact extraction task, we evaluated the accuracy of both the facts extracted from unstructured biomedical text and the supporting evidence for each gene-fact statement. Manual evaluation was performed on a random subset of ten publications from our set (see Methods section), for a total of 165 gene-fact pairs. For each extracted gene-fact pair, the following were assessed: (i) precision, defined as whether the extracted statement was correct and supported by verbatim text extracted from the source paper (ii) recall, defined as whether relevant gene facts or data types were retrieved by the model. We have also computed the F1-score at the overall paper and data-type levels.

For the gene summary generation task, we evaluated the ability of GeneScribe to produce concise and accurate textual descriptions from the structured facts extracted in the first stage. The generated summaries were manually assessed by comparing the previous Alliance summary and the updated summary. The evaluation calculated summary precision, recall and F1 score, assessing whether the generated summary faithfully reflected the extracted facts without introducing unsupported facts. We have also considered readability, defined as whether the summary was clear, coherent, and suitable as a gene summary.

## Results

### Gene fact extraction results

Across 83 selected publications (see Methods Section), the pipeline extracted gene-fact statements related to 212 genes, 509 biological processes, 212 molecular functions, 140 gene interactions, 210 pathways 186 gene expressions, and 43 disease associations (Figure. 2). These results indicate that the structured prompting framework can capture diverse categories of biologically relevant information directly from unstructured scientific literature.

**Figure 2.**
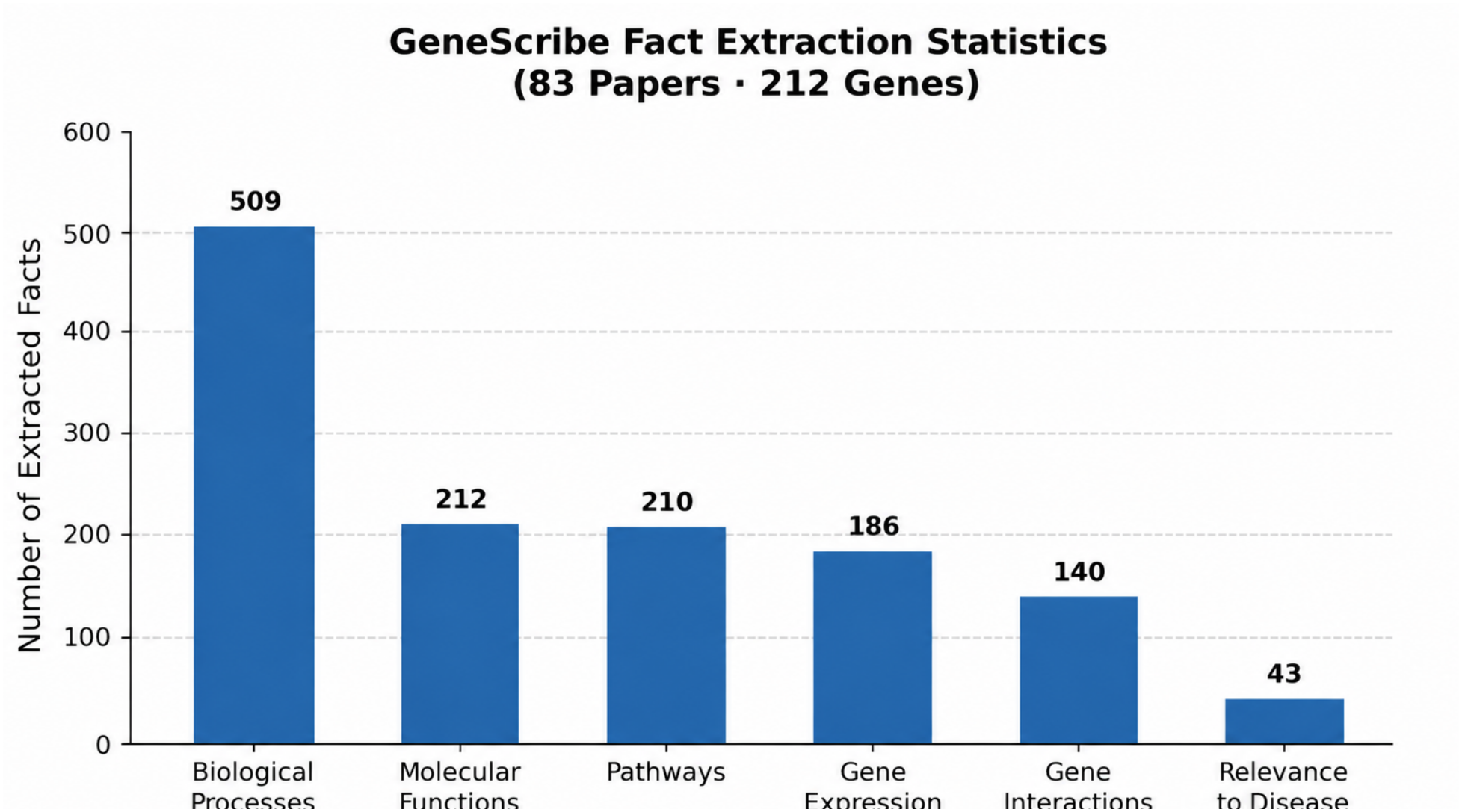
GeneScribe fact extraction statistics

159 extracted gene-fact pairs were judged to be correct, while 6 were marked incorrect, corresponding to a precision of 96.4%. These results indicate that the schema-constrained prompting strategy effectively limits unsupported or hallucinated biological statements. Across the reviewed publications, 31 relevant gene-fact pairs were identified as missed by the extraction pipeline, corresponding to a recall of 83.7% and an overall F1-score of 89.6%. Missed facts most frequently involved pathway associations, localization information, and disease-relevance statements.

We further analyzed extraction performance at the individual paper level. As shown in Figure. 3, GeneScribe achieved consistently high precision across most publications, with seven of the ten evaluated papers reaching 100% precision, indicating that the extracted facts were highly reliable. Recall showed greater variation across papers, reflecting differences in the complexity and explicitness of the reported biological findings. Overall, GeneScribe achieved average precision, recall, and F1-scores of 92.2 ± 5.1%, 78.9 ± 4.3%, and 84.4 ± 4.2% (mean ± SE across 10 papers), respectively, demonstrating robust fact extraction performance across diverse biomedical publications. One paper, P9, showed a lower score (F1-score =54.5%). We attribute this issue to these factors: (i) the model missed identifying one of the genes described in the paper and hence no related facts were extracted for this gene (ii) inaccurate molecular function, biological process and sub-cellular localization attributions to another gene identified in the paper, possibly due to language in the paper that made it difficult for the model to clearly identify these datatypes for this gene as defined in the prompt. The latter is often a common issue in biocuration even for manual curation tasks which is resolved by the curator’s training and expertise. We think that giving the model access to more comprehensive curation manuals and best practices will improve performance in future work.

**Figure 3.**
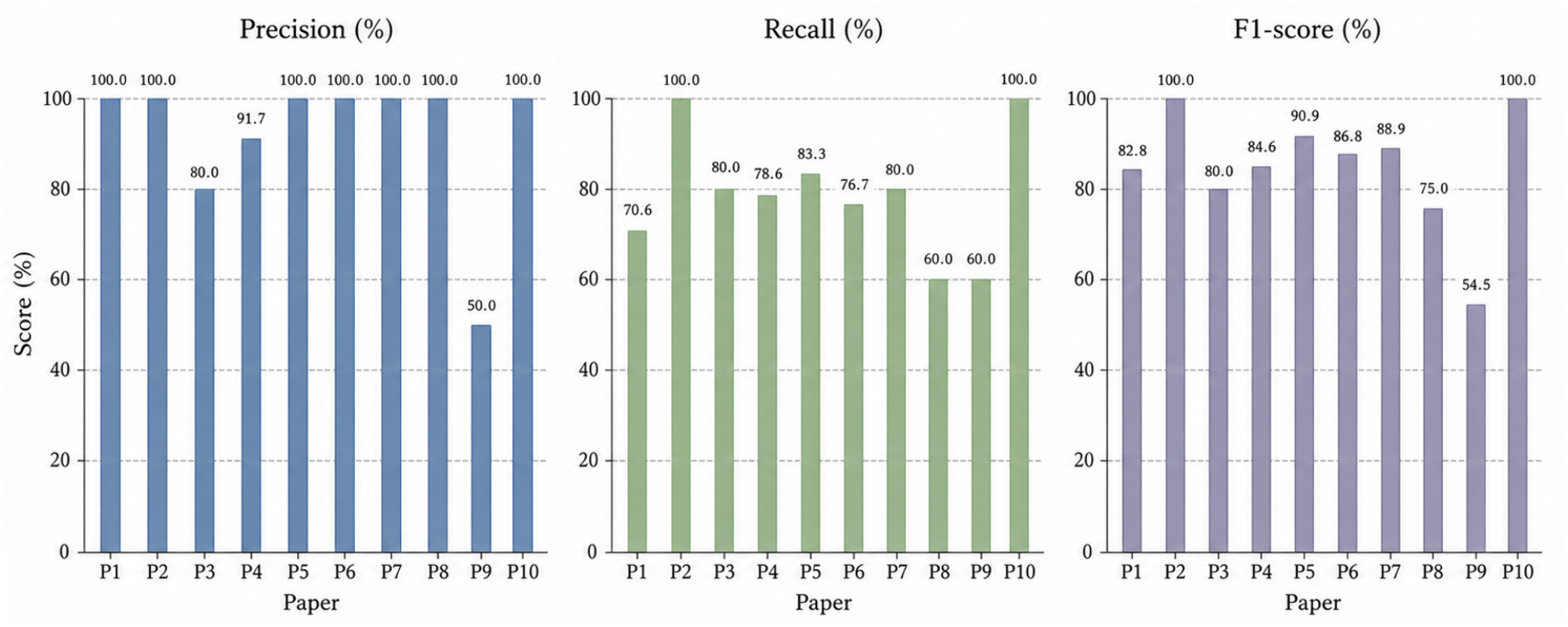
Paper-level fact extraction performance of GeneScribe across ten manually evaluated publications. Precision, recall, and F1-score were computed independently for each paper based on manually curated gene-fact annotations.

Curators additionally evaluated whether the evidence sentences returned by the model were sufficient to support the extracted facts and documented recurring extraction errors. The most common issue involved was related to the extraction of evidence sentences by the LLM (total of 6 cases). There were 3 cases where there was evidence attribution mismatch, where the extracted fact itself was correct but the retrieved evidence sentence was not. In 3 cases, both the fact and the evidence sentence extracted was incorrect. Another issue involved pathway interpretation errors, where the model inferred that a gene was part of a signaling pathway when the paper only showed that the gene affected or regulated that pathway.

Overall, these results demonstrate that GeneScribe achieves high factual precision while producing structured, evidence-grounded outputs suitable for downstream biocuration workflows and automated gene summary generation.

### Gene summary generation results

We evaluated whether GeneScribe could synthesize concise and readable gene summaries from the extracted literature-derived facts. The 10 papers we randomly selected for our test set contained mentions of 56 genes in the titles or abstract. We evaluated the gene summaries for these genes, 46 genes with existing summaries and 10 genes with updated summaries. We compared the previous Alliance summary (when present), the extracted statements from papers, and the GeneScribe-generated summary. All 56 summaries were manually evaluated by a curator. Of these, 54 were judged correct and 2 were judged incorrect, corresponding to an overall summary-generation accuracy of 96.43%. Examination of the incorrect summaries revealed that the challenge was not related to hallucinations in generated text, but the faithful representation of the facts (whether predicted or experimental) and the preservation of all of the content of the previously existing Alliance summary.

For readability, 53 summaries were judged readable and 3 marked as having reduced readability, corresponding to a readability rate of 94.64%. These results suggest that GeneScribe generally produces biologically meaningful and readable summaries. For example, for *cdc-42*, the previous Alliance summary already included molecular function, biological processes, localization, expression, disease relevance, and human orthology information. GeneScribe successfully incorporated the newly extracted facts related to cell polarity, PAR protein polarity, actomyosin cytoskeleton regulation, interaction with MRCK-1, Rho protein signal transduction, and cell cortex localization, in order to build a new and readable gene summary. In a second example, the gene *acdh-1* was correctly described as “involved in the biological process propionate catabolic process” and “enabling the molecular function acyl-CoA dehydrogenase activity,” both of which align with curated experimental evidence.

Considering the results presented in this section, we think that the performance of GeneScribe is adequate for both fact extraction and gene summary generation tasks and is therefore suitable for a future inclusion into the biocuration workflow at the Alliance.

## Discussion

GeneScribe offers a two-step process for generating gene summaries where gene facts are extracted by data type along with the provenance and then a second step where summaries are built based on the extracted facts. For genes that lack a summary at the Alliance, GeneScribe generates summaries from newly extracted facts directly from the literature without waiting for curation to happen first. For genes that already have a summary, GeneScribe preserves the existing Alliance summary which is based on curated data while integrating newly extracted literature-derived findings. This strategy allows gene summaries to evolve incrementally as new publications become available, minimizes unnecessary gene summary changes, and maintains consistency across Alliance gene summaries. By enforcing a set of structured guidelines (sentence templates, data type definitions, etc.) the model produces concise yet informative gene summaries that align stylistically and scientifically with existing automated summaries. Full provenance is maintained through evidence in the form of extracted text from the paper, so that facts can be traced back to the source publication, and are available for reuse in downstream biocuration workflows.

In conclusion, GeneScribe provides an effective method to generate gene summaries based on the relevant literature without relying solely on previously curated data and can scale to match the rate at which gene-related data is published. This aligns with the current Alliance goals of improving automation using AI in the biocuration workflow (Arnaboldi et al., 2026).

## Data availability statement

The complete prompts given to the LLM for the different tasks along with the code is available at this Alliance GitHub repository (http://github.com/alliance-genome/agr_genescribe). All data used in this paper were obtained from the Alliance version 8.1.0.

## Acknowledgement

Generative AI was used as an integral component of this research for automated biological fact extraction from scientific literature and gene function summary generation. OpenAI’s GPT-o1 model was primarily used, with GPT-4o and other GPT models evaluated for comparison. Claude AI and ChatGPT were used to assist with polishing the manuscript. After using these tools, the authors reviewed and edited the content as needed and take full responsibility for the accuracy and integrity of the published article.

## Funding

The Alliance is supported by NIH NHGRI and NHLBI: U24HG010859 and WormBase is supported by NIH NHGRI U24HG002223.

## Conflict of Interest

The authors declare that they have no conflict of interest.

